# An Evolutionary Conserved Multi-Stress Sensory Histidine Kinase NblS Associates With Photosystem II Proteins And Responds To Its Redox Status In The Cyanobacterium *Synechococcus elongatus* PCC 7942

**DOI:** 10.1101/2025.01.21.633742

**Authors:** Tatsuhiro Tsurumaki, Kan Tanaka

**Author notes:** **Corresponding Authors:** T. Tsurumaki, Centre Algatech, Institute of Microbiology of the Czech Academy of Sciences, Novohradská 237, Třeboň 379 01, Czechia. Subject Areas: (2) environmental and stress responses, (3) regulation of gene expression.

## Abstract

Responding to stress caused by various environmental changes is essential for living organisms. In cyanobacteria that perform oxygenic photosynthesis, the highly conserved histidine kinase Hik33/NblS homologs respond to diverse stressors such as high light, low temperature, high salts, high osmolarity, and reactive oxygen species. However, how this unique protein kinase responds to such divergent stresses remains unknown. This study has focused on the underlying stress sensing mechanism of NblS in *Synechococcus elongatus* PCC 7942. First, the sensory response by NblS was analyzed *in vivo* by monitoring the NblS-regulated *hliA* transcript accumulation with treatment of various benzoquinone reagents known as photosystem II (PSII) electron acceptors. It was found that molecular responses induced by various stresses were diminished in the presence of 2,6-dichloro-1,4-benzoquinone, which accepts electrons specifically from the PSII-bound plastoquinone Q_B_. Cell fractionation analysis indicated that NblS was localized in the thylakoid membrane, which was consistent with its predicted membrane-spanning structure. In the thylakoid membrane, NblS was found in approximately 400 kDa and 800 kDa unknown complexes in clear native PAGE (CN-PAGE). Immunoprecipitation analysis of the cross-linked thylakoid membrane revealed that NblS is associated with D2 and CP47 proteins but not with CP43 protein, and thus it was suggested that dimeric NblS is associated with RC47-like complex, an assembly intermediate complex of PSII, as [RC47like-NblS_2_] (370 kDa) or [RC47like-NblS_2_]_2_ (740 kDa). We propose that the redox status of an RC47-bound plastoquinone molecule is a cue for the NblS response.

## Introduction

<u>T</u>wo-<u>C</u>omponent-<u>S</u>ystem (TCS) is a major module for sensing and responding to environmental changes in bacteria. TCS is composed of a sensory histidine kinase (HK) and a cognate response regulator (RR), which is typically a transcription factor, where signaling occurs as the phospho-transfer reaction from HK to RR resulting in the control of gene expression at the transcriptional level (Laub and Goulian, 2007). Cyanobacteria are the only bacterial phylum that performs oxygenic photosynthesis. Thus, Cyanobacteria have played a major role in developing and maintaining the Earth’s environment. Among many TCSs that have been found in cyanobacteria, it has been noticed that a set of TCS components are specifically conserved in cyanobacteria (Ashby and Houmard, 2006), suggesting a strong relationship with oxygenic photosynthesis. In this study, we focused on one of such TCS components, Hik33/NblS homologs.

Hik33/NblS homolog was first identified by genetic analysis in the unicellular cyanobacterium *Synechocystis* sp. PCC 6803 (hereafter *Synechocystis*), where missense mutations resulting in resistance to several herbicides were identified in the gene named *dspA* (<u>d</u>rug sensory <u>p</u>rotein <u>A</u>) (Bartsevich and Shestakov, 1995). Subsequently, the first cyanobacterial whole genome project in *Synechocystis* identified 43 genes for HK (Mizuno et al. 1996), and the exhaustive functional analyses using insertional knock-down mutants and microarray analyses provided evidence that Hik33, a synonym of DspA, is responsive to multiple stresses such as high light (HL), cold, high salt, high osmolality, and H_2_O_2_ (Kanesaki et al. 2007; Marin et al. 2003; Mikami et al. 2002; Suzuki et al. 2001; Tu et al. 2004). Photosynthesis-related genes including high light-inducible protein genes (*hli* genes) belonging to the CAB/ELIP/HLIP superfamily and genes of both photosystems (PSII and PSI) components were found to be under the control of Hik33, indicating tight relationship with the photosynthetic light reactions (Hsiao et al. 2004).

A gene encoding Hik33 homolog was also identified in *Synechococcus elongatus* PCC 7942 (hereafter *Synechococcus*) by genetic analysis of a mutant showing decreased induction of *hliA*, one of the *hli* genes, during HL and UV-A irradiation (Van Waasbergen et al. 2002). Because this mutant showed a nonbleaching phenotype: defective phycobilisome degradation under nitrogen or sulfur depletion, the mutated HK gene was named *nblS* after the <u>n</u>on<u>bl</u>eaching sensor. Introduction of *Synechococcus nblS* gene into the *Synechocystis hik33* mutant deficient in the HL-inducibility of *hli* genes complemented the phenotype (Hsiao et al. 2004), thus indicating that these two genes are functionally orthologous.

*In vitro* analysis using recombinant proteins showed that NblS has both kinase and phosphatase activities targeting an RR named RpaB (López-Redondo et al. 2010). Under physiological growth light conditions, RpaB is phosphorylated and binds to a conserved sequence element, HLR1. Under HL conditions, RpaB is rapidly dephosphorylated and dissociates from DNA (Hanaoka and Tanaka, 2008; Kato et al. 2022; Moronta-Barrios et al. 2012). Furthermore, the dephosphorylated RpaB level *in vivo* was well correlated with the light intensity to which the cells were exposed (Yasuda et al. 2020). These observations suggested that Hik33/NblS homologs function as a kinase under physiological growth conditions and works as a phosphatase upon endowed stresses. Recently, reduction of a photosynthetic electron transport component, possibly Q_A_ of PSII or plastoquinone (PQ) pool, was proposed as the signal to induce Hik33/NblS dependent response (Kato et al. 2022; Mironov et al. 2019). However, the NblS-sensory mechanism still remains elusive.

In the present study, we investigated how the NblS activity is regulated in *Synechococcus*. Our transcriptional analysis revealed that NblS responds to a series of stressors the same as *Synechocystis* Hik33. Furthermore, all of the responses were quenched by benzoquinone reagents that oxidize Q_B_ in PSII. Then, we studied the intercellular localization and NblS protein complex by constructing FLAG-tagged NblS. We showed that NblS localizes in the thylakoid membrane and binds to PSII subunits, CP47 and D2. These results indicate that NblS is a stress sensor of PSII. Our phylogenetic analysis revealed that Hik33/NblS orthologs are composing a monophyletic clade distinct from other HKs, suggesting the conserved stress sensing mechanism among orthologs.

## Results

### NblS-dependent HL response corresponds to the reduction of PSII-bound Q_A_

To examine the hypothesis that NblS is responsive to the reduction of Q_A_ or PQ pool (Kato et al. 2022; Mironov et al. 2019), we studied whether the oxidation of PSII would quench the NblS signaling. Benzoquinone derivatives have been used as electron acceptors to modulate the electron transport around PSII (Wiwczar and Brudvig, 2017). We used *hliA* transcript accumulation as an index to monitor the NblS response (Yasuda et al. 2020). The NblS-dependent HL response was examined in the presence of 1,4-benzoquinone (BQ), 2,6-dichloro-1,4-benzoquinone (DCBQ), 2,6-dimethyl-1,4-benzoquinone (DMBQ) or tetramethyl-1,4-benzoquinone (duroquinone: DQ), where efficiency of electron transfer from Q_A_, Q_B_ or PQ pool differs depending on the benzoquinone molecule species (Fig. 1A). As the results, the *hliA* transcript accumulation was completely inhibited by BQ or DCBQ, while alleviated by DMBQ or DQ (Fig. 1B). It is known that DCBQ has the highest affinity to Q_B_ among these acceptors (Fu et al. 2017; Satoh et al. 1992). Thus, the differential effects of benzoquinone derivatives suggested that the reduced state of PSII but not PQ pool correlated with the NblS signaling. This hypothesis was supported by the observation that a non-quinone type electron acceptor from Q_A_, silicomolybdate (Schansker and van Rensen, 1993) decreased the *hliA* transcript accumulation in a concentration dependent manner (Fig. S1A). The difference of quenching effect between DCBQ and silicomolybdate could be due to their accessibility to PSII in intact cells, where the DCBQ has a far smaller molecular weight (MW) than silicomolybdate, 177 and 1823, respectively. In contrast to the *hliA* gene regulation, transcription of the PSI reaction center protein genes *psaAB* is positively regulated by RpaB under growth light and downregulated upon shift to HL (Kato et al. 2022). We examined the DCBQ effect under HL conditions and found that the *psaAB* transcript did not decrease in the presence of DCBQ under HL (Fig. S1B).

**Figure 1.**
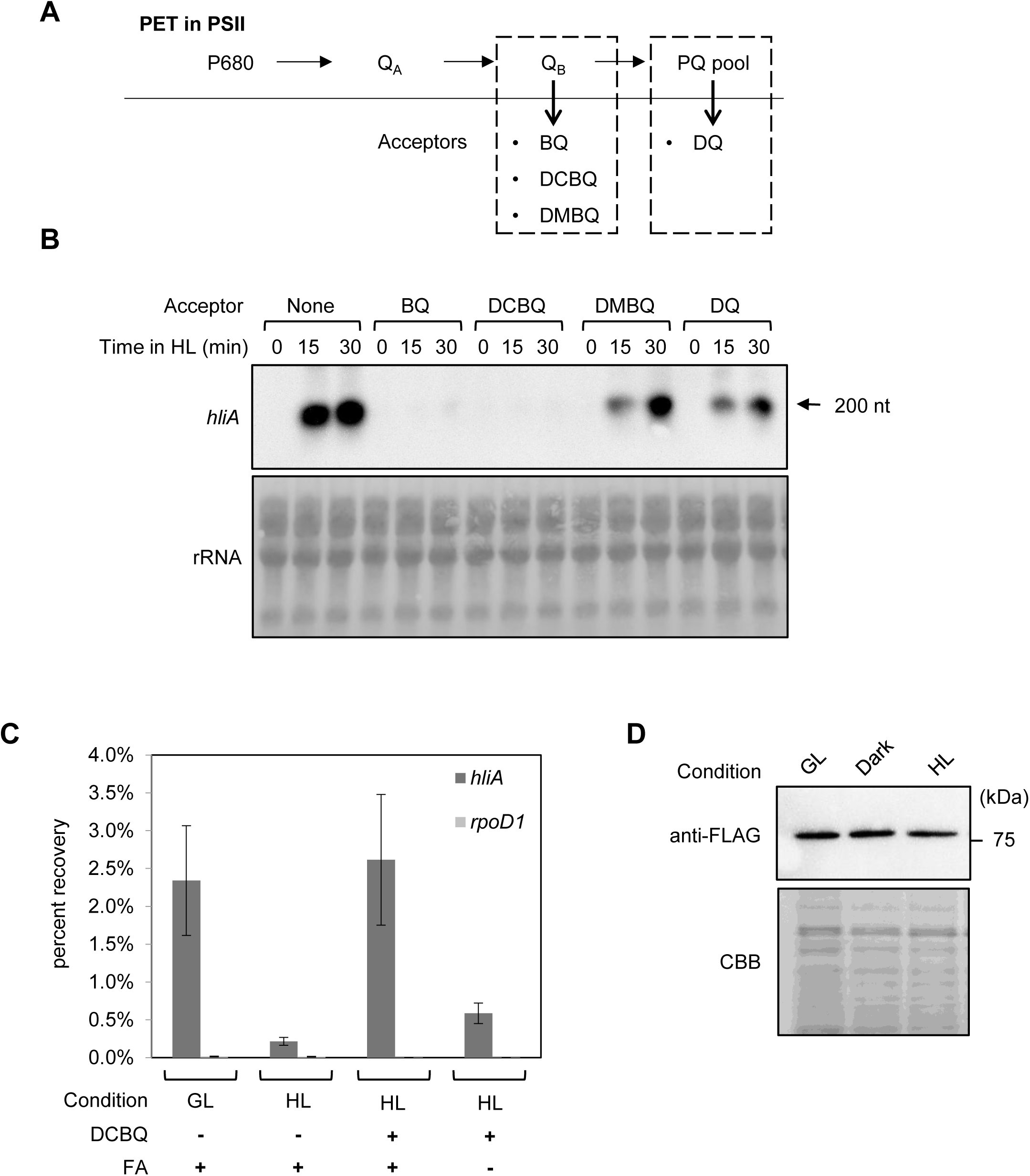
Quenching analysis of HL stress response by PSII alternative electron acceptors. (**A**) The cartoon of simplified photosynthetic electron transport (PET) in PSII and action sites of each electron acceptors. (**B**) the result of Northern blot for *hliA* transcript under HL condition with several electron acceptors. Each 100 µM acceptors were added to *Synechococcus* WT culture respectively for 5 min before HL irradiation. rRNA is the image of Methylene Blue stained membrane as the loading control. 3 µg of total RNA were loaded in each lane. (**C**) The result of ChIP-qPCR of RBF strain under HL condition with DCBQ. 100 µM DCBQ were added to *Synechococcus* RBF culture and incubated for 5 min under growth light (GL). Then, HL treatment was done for 5 min. FA indicates the sample treatments with (+) or without (-) formaldehyde for ChIP. Data are means ± SD from three independent experiments. (**D**) The expression analysis of NblS-FLAG under different light conditions by immunoblot. *Synechococcus* NF strain cultivated under GL were transferred to HL or Dark for 5 min, respectively. The cell lysates corresponding to 0.5 µg chlorophyll were used for SDS-PAGE and Immunoblot. NblS-FLAG was detected by anti-FLAG antibody. CBB is the image of Coomassie Brilliant Blue stained gel as the loading control.

The phosphorylated RpaB protein binds to HLR1 elements in the *hliA* promoter region to repress the transcription under physiological conditions. RpaB is desphosphorylated under HL in an NblS-dependent manner, resulting in its dissociation from the promoter and the activation of the *hliA* promoter. Given that DCBQ abolished the *hliA* promoter activation under HL, we expected that RpaB protein remained bound to the promoter region. Thus, chromatin immunoprecipitation-qPCR analysis was performed using a FLAG-tagged RpaB expressing strain as previously reported (Hanaoka and Tanaka, 2008). The results confirmed that the RpaB dissociation from the promoter region under HL was inhibited by DCBQ (Fig. 1C).

Furthermore, we constructed a FLAG-tagged NblS expressing strain (hereafter NF strain) to monitor the NblS protein accumulation during the HL stress response. The result revealed that the NblS protein amount remains constant under the examined conditions and thus the signaling events were not accompanied by the NblS protein level changes (Fig. 1D).

### Lines of evidence supporting the reduced Q_A_ hypothesis of the NblS sensing

An herbicide 3-(3,4-dichlorophenyl)-1,1-dimethylurea (DCMU) binds to the Q_B_ site of PSII and blocks the electron transport from Q_A_ to Q_B_. It was previously found that DCMU treatment partially mimics the HL response of gene expression in both *Synechocystis* and *Synechococcus* (Hihara et al. 2001; Salem and Van Waasbergen, 2004), and we examined the effect of DCMU in the presence of DCBQ. As the result, DCMU induced the *hliA* transcription and the response was quenched by DCBQ (Fig. 2), which is consistent with the Q_A_ sensing model. We also showed the occurrence of responses under cold, H_2_O_2_, salt, and osmotic stress conditions in *Synechococcus* as found in *Synechocystis*, and all of these responses were abolished in the presence of DCBQ (Fig. 2). These results support the previous suggestion of the reduced Q_A_ hypothesis of the NblS sensing.

**Figure 2.**
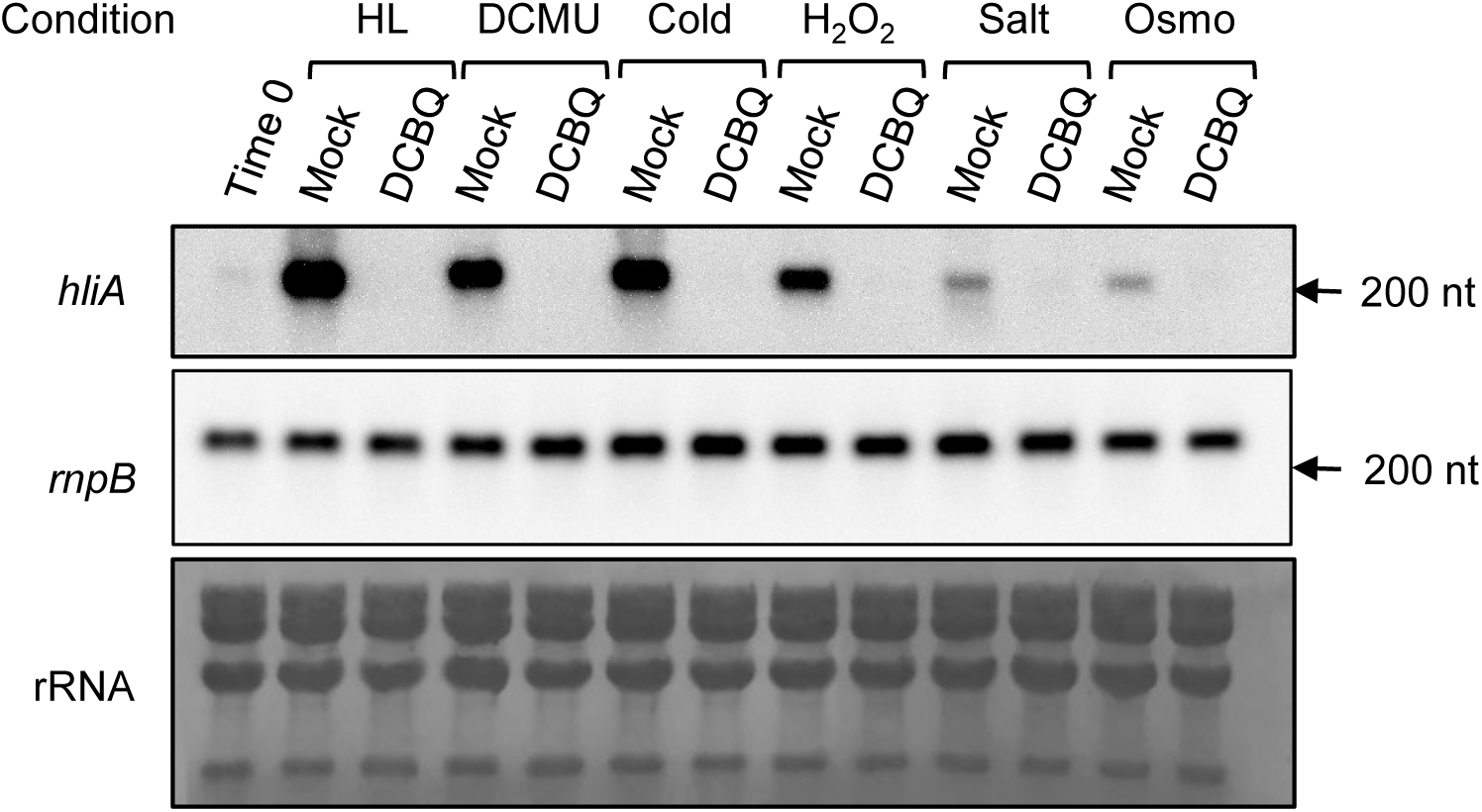
Northern blot analysis of *hliA* transcript under various stress conditions with DCBQ. One culture of *Synechococcus* WT grown under normal condition were separated and transferred to stressed condition for 30 min as HL, DCMU: 10 µM of DCMU, Cold: 14°C, H_2_O_2_: 0.25 mM H_2_O_2_, Salt: 0.5 MNaCl and Osmo: 0.5 M sorbitol respectively. Mock and DCBQ indicate the samples treated with 0.1% ethanol or 100 µM DCBQ for 5 min before transferring to stress conditions respectively. Time 0 indicates the sample before any treatments. *hliA* and *rnpB* were detected by each specific probes. *rnpB* is the internal control. rRNA is the image of Methylene Blue stained membrane as the loading control. 3 µg of total RNA were loaded in each lane.

### NblS is localized in the thylakoid membrane fraction

In *Synechocystis*, Hik33 was shown to be integrated in the thylakoid membrane (Wang et al. 2022), however, there is no relevant information in *Synechococcus*. In silico analysis of the NblS secondary structure predicted that there are two transmembrane segments in the N-terminal region as found in Hik33 (Fig. 3A), which suggested that NblS is also a similarly membrane-integrated protein. To identify the cellular localization of NblS, whole cell lysate was separated by sucrose density gradient centrifugation (Schottkowski et al. 2009) and NblS protein was analyzed by immunoblot. NblS was detected from fractions IV and V, which were enriched with thylakoid membrane protein D1, the PSII reaction center protein (Fig. 3B). Meanwhile, the NblS protein was not found in fraction II which was enriched with the plasma membrane. It has been known that thylakoid membrane is heterogeneously composed with different protein distributions (Huokko et al. 2021). Thus, the fraction V including the NblS protein was further fractionated by the second-round centrifugation as described previously in *Synechocystis* (Schottkowski et al. 2009). As a result, NblS was co-fractionated with D1 (Fig. 3B), indicating that NblS signaling events are occurring on the PSII-containing thylakoid membrane.

**Figure 3.**
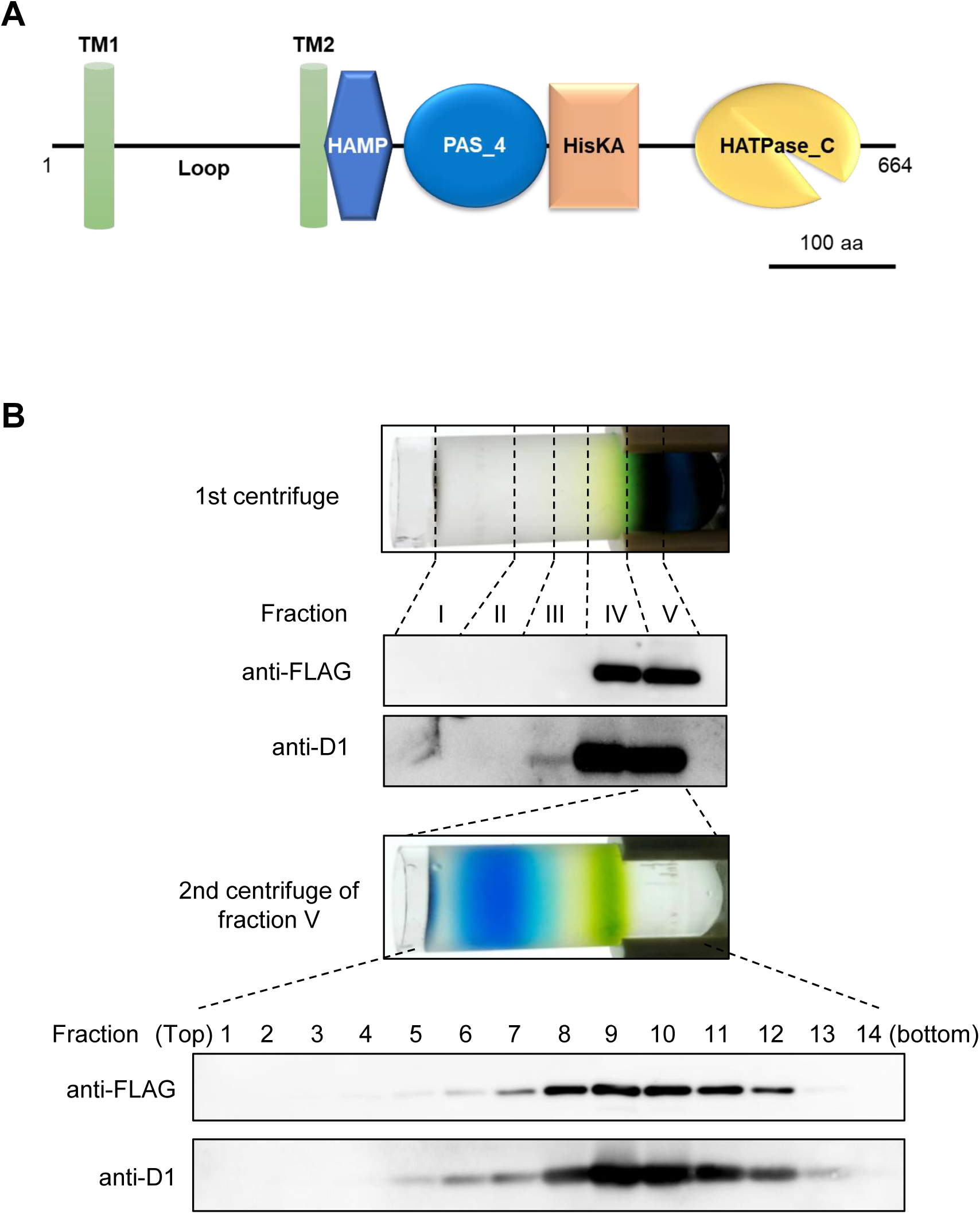
Analysis of NblS-FLAG localization by cell fractionation. (**A**) The secondary structure of NblS (Pfam: Q8RQ68). TM and Loop indicate transmembrane regions and putative luminal loop, respectively. Other annotations are according to Pfam. (**B**) Fractionation of cell lysate from NF strain. Top: the picture of the centrifuge tube after first centrifugation. Positions of five fractions are indicated by broken lines. Middle: Fraction V was used to second centrifugation and the picture of centrifuge tube after centrifugation is shown. From top to bottom of the tube, 14 fractions with equal volume were collected. NblS-FLAG and D1 were detected by specific antibodies.

### NblS is associated with PSII subunit proteins

To examine whether NblS is associated with other proteins, protein complexes in the thylakoid membrane fraction were separated by clear native-PAGE (CN-PAGE) as described previously (Koskela et al. 2020). The NblS protein was detected at two MW positions larger than the 150 kDa of the expected NblS dimer (hereafter NblS_2_). In addition, the pattern of NblS complexes was stable after the HL treatment of isolated thylakoid membrane (Fig. 4A).

**Figure 4.**
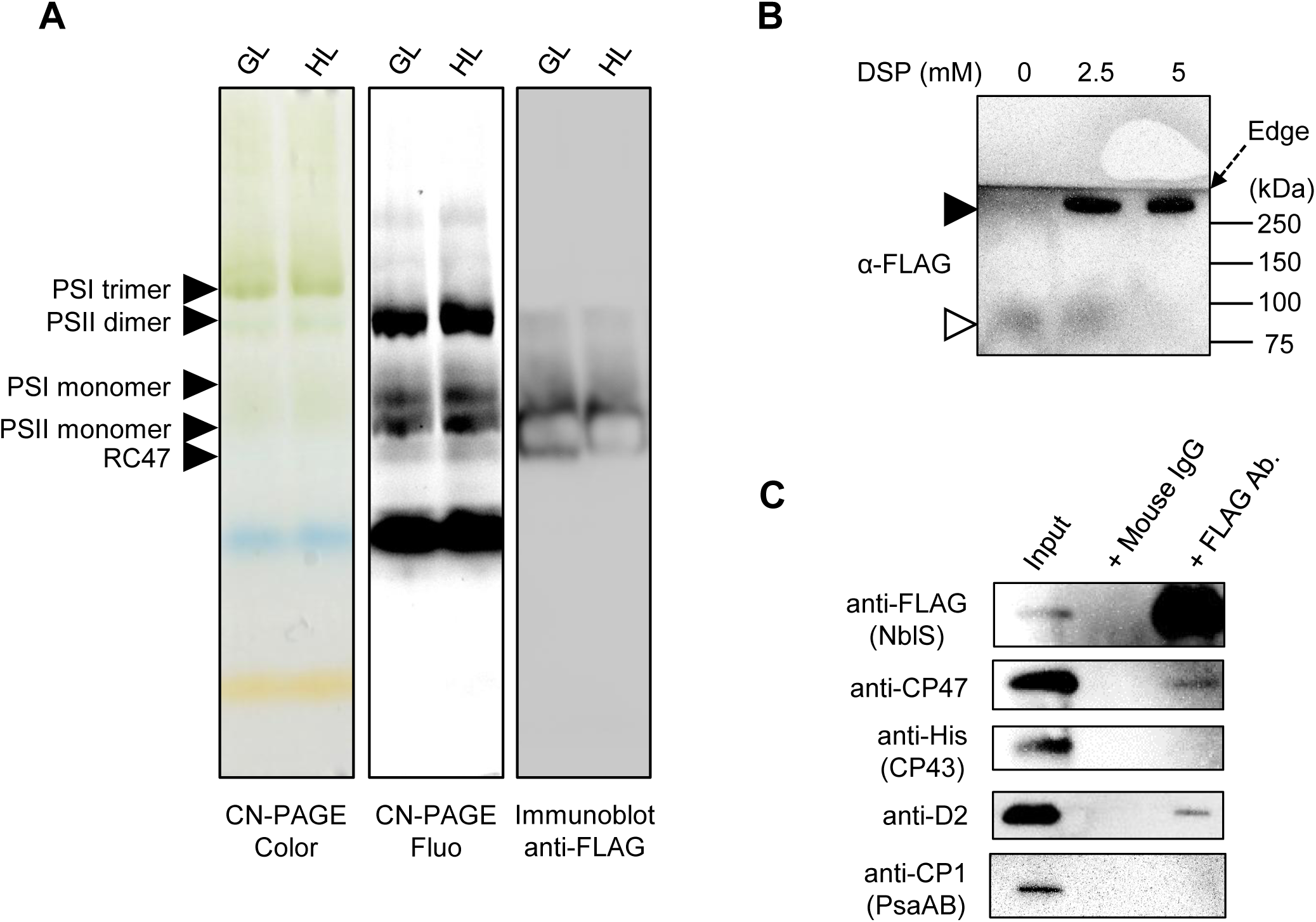
Detection and purification of NblS protein complexes in the thylakoid membrane. (**A**) Thylakoid membrane protein complexes of NF43H strain were separated by CN-PAGE. CN-PAGE Color is the RGB gel picture, CN-PAGE Fluo is the chlorophyll fluorescence gel image excited at 505±25 nm and detected with a red filter. The positions corresponding to each photosystem complexes are indicated according to the previous work (Koskela et al. 2020). The position of NblS-FLAG detected by immunoblot is shown in Immunoblot anti-FLAG. HL 1 min indicates the thylakoid membrane sample exposed to 2,000 μmol m^-2^ s^-1^ white LED light for 1 min prior to solubilization. (**B**) Immunoblot of DSP cross-linked thylakoid membrane after non-reduced SDS-PAGE. DSP treatments were done with indicated concentrations respectively. White arrowhead and black arrowhead indicate non-cross-linked and cross-linked NblS-FLAG, respectively. The image includes the upper edge of the PVDF membrane (Edge). (**C**) Immunoblot of the protein sample after FLAG immunoprecipitation with DSP cross-link. SDS-PAGE was done after cleavage of DSP cross-link by DTT. Each protein was detected by indicated antibodies. Input lane indicates the thylakoid membrane of NF43H strain corresponding to 0.5 µg chlorophyll. +Mouse IgG lane and +FLAG Ab. lane indicate the precipitated samples by nonspecific and specific antibody, respectively.

To purify the putative NblS complex from the solubilized thylakoid membrane fraction, immunoprecipitation was performed using anti-FLAG antibody (Fig. S2A). However, only the NblS protein itself was detected. Therefore, the NblS protein was first cross-linked by a membrane-permeable protein cross-linker dithiobis (succinimidyl propionate) (DSP) in the cell as previously reported (Liu et al. 2013). After cross-linking, the migration pattern of NblS protein complexes in the immunoblot after CN-PAGE was the same as in non-cross-linked sample (Fig. S2B). In addition, a protein complex containing NblS was detected at a position larger than 250 kDa by non-reduced SDS-PAGE and immunoblot, indicating that the protein complex containing NblS was successfully stabilized under cross-linked condition (Fig. 4B).

Taking into account that the occurrence of NblS signaling events correlates with the Q_A_ redox status in PSII (Fig. 2), the interaction with PSII or PSII-related complex was considered. To examine this, the *psbC* gene encoding the PSII core antennae protein CP43 of the NF strain was hexahistidine (His_6_)-tagged as described (Sugiura and Inoue, 1999), and the resultant strain NF43H was used for subsequent analysis. In this study, we failed to introduce His_6_-tag in another antennae protein CP47, which also has been used to analyze PSII complexes in *Synechocystis* (Bricker et al. 1998). Probably, it was due to the essentiality of the operon (Gene ID: *Synpcc7942_0694* to *Synpcc7942_0697*) including PsbT encoding gene *psbB* (*Synpcc7942_0694*) (Rubin et al. 2015; Vijayan et al. 2011), while the operon is not composed in *Synechocystis*.

The cross-linked thylakoid membrane fraction from NF43H was subjected to the pull-down procedure for the FLAG-tagged NblS. Precipitated proteins were reduced to cleave the disulfide bond from DSP-crosslinks and subjected to immunoblot analysis with antibodies against PSII subunit proteins. As a result, we identified the PSII reaction center D2 protein and the core antenna CP47 protein in the NblS associating complex (Fig. 4C), while another core antenna CP43 protein (CP43-His_6_) was not detected. Thus, we suggested that NblS is associated with the CP43-less PSII-related complex. Consistently, a pull-down experiment with the CP43-His_6_ by Ni^2+^ resin did not precipitate NblS while pull-down of PSII was confirmed by the detection of the D2 protein (Fig. S2C). In both FLAG-tag and His-tag pull-down experiments, PSI reaction center protein PsaAB was not detected in the precipitate (Fig. 4C, S2C).

To support our protein cross-linked co-immunoprecipitation result, we evaluated the NblS_2_-CP47-D2 protein complex by modeling through Alphafold3 (Abramson et al. 2024). Since protein structures of CP47 and D2 have been well solved and accumulated in protein data bank (PDB) database (Bekker et al. 2022), both of them showed high pLDDT score (Fig. S3B). On the contrary, a relatively lower pLDDT score to NblS_2_ was obtained due to the low number of datasets of HK structures. The modeled complex showed that NblS_2_ locates next to the third helix of CP47 (Fig. S3B). In this study, we used a protein cross-linker DSP which reacts with primary amine in lysine (Lys) residue, then we highlighted them in the structure (Fig. S3B). As the result, K137 of CP47 and K24 of NblS showed the closest distance 9.828 Å which is within 12 Å of the spacer arm of DSP. Thus, it suggests that these Lys residues are most likely cross-linked in the protein complex.

### Phylogenetic analysis of NblS indicates conservation and generality of the function

To estimate how far the Hik33/NblS orthologs are conserved among lineages, we constructed a phylogenetic tree with HKs which show significant similarity to Hik33/NblS. It has been known that chloroplast genomes in red algae retain the Hik33/NblS orthologs named Ycf26 (Puthiyaveetil and Allen, 2009), so we extended our analysis to eukaryotic plastids as well as cyanobacteria. Since the N-terminal sequence of NblS until TM2 (Fig. 3A) is unique within cyanobacteria proteomes, the region of functional domains in NblS, e.g. HAMP-PAS-HisKA-HATPase (Fig. 3A), was used as the query for the protein similarity search by BLASTP (Altschul et al. 1990). To keep the integrity of NblS function, the result of BLAST was filtered to the query coverage more than 90%. Furthermore, taking into account the gene duplications and domain shuffling in cyanobacterial HKs (Ashby and Houmard, 2006; Narikawa et al. 2004), not the full length sequences but only aligned sequences of hit proteins were retrieved and used for phylogenetic analysis. Regarding the conservation of Hik33/NblS homologs, we found them from 305 out of 308 cyanobacteria species in our genome database (Supplementary Dataset S1). Only three species were not hit by BLASTP although DNA sequences with coverage more than 80% and quite low E-value were found in all of them by TBLASTN (Supplementary Dataset S1 and S2). It most probably indicates that Hik33/NblS homologs are conserved in all cyanobacteria.

As stated above, partial protein sequences were employed to the phylogenetic tree, however, the domain structure from the full length of them were plotted in the tree to visualize their diversity (Fig. 5). Even though some HKs have the same domain composition as Hik33/NblS (e.g., TM-HAMP-PAS-HisKA-HATPase without the additional domain), Hik33/NblS orthologs form an own clade that differs from other HKs (Fig. 5). Interestingly, all Hik33/NblS orthologs have no additional domains in their own structures throughout all organisms. All cyanobacterial orthologs showed more than 50% identity to the NblS query sequence, while eukaryotic orthologs showed more than 35% identity. The other HKs showed identity less than 32%. It is also noteworthy that Hik33/NblS orthologs branched from Melainabacteria, known as the non-photosynthetic sister clade of cyanobacteria (Sánchez-Baracaldo and Cardona, 2020), as the root. Next of Melainabacteria, thylakoid-less cyanobacteria *Gloeobacter* and *Anthocerotibacter* were located as the root of photosynthetic cyanobacteria (Rahmatpour et al. 2021). In agreement with the previous report (Ponce-Toledo et al. 2017), eukaryotic clade branched from cyanobacterium *Gloeomargarita lithophora*, which was previously proposed as the last common ancestor of chloroplast by phylogenetics, to a Glaucophyte *Glaucocystis incrassate* and to a Rhodophyte. Then, Hik33/NblS orthologs from marine *Synechococcus*/*Prochlorococcus* and heterocystous cyanobacteria formed specific clades from other cyanobacteria. This tree topology was consistent even when full length of NblS orthologs was used for the analysis (Fig. S5). Thus, the functional domains of Hik33/NblS comprise the distinct and general function different from other HKs.

**Figure 5.**
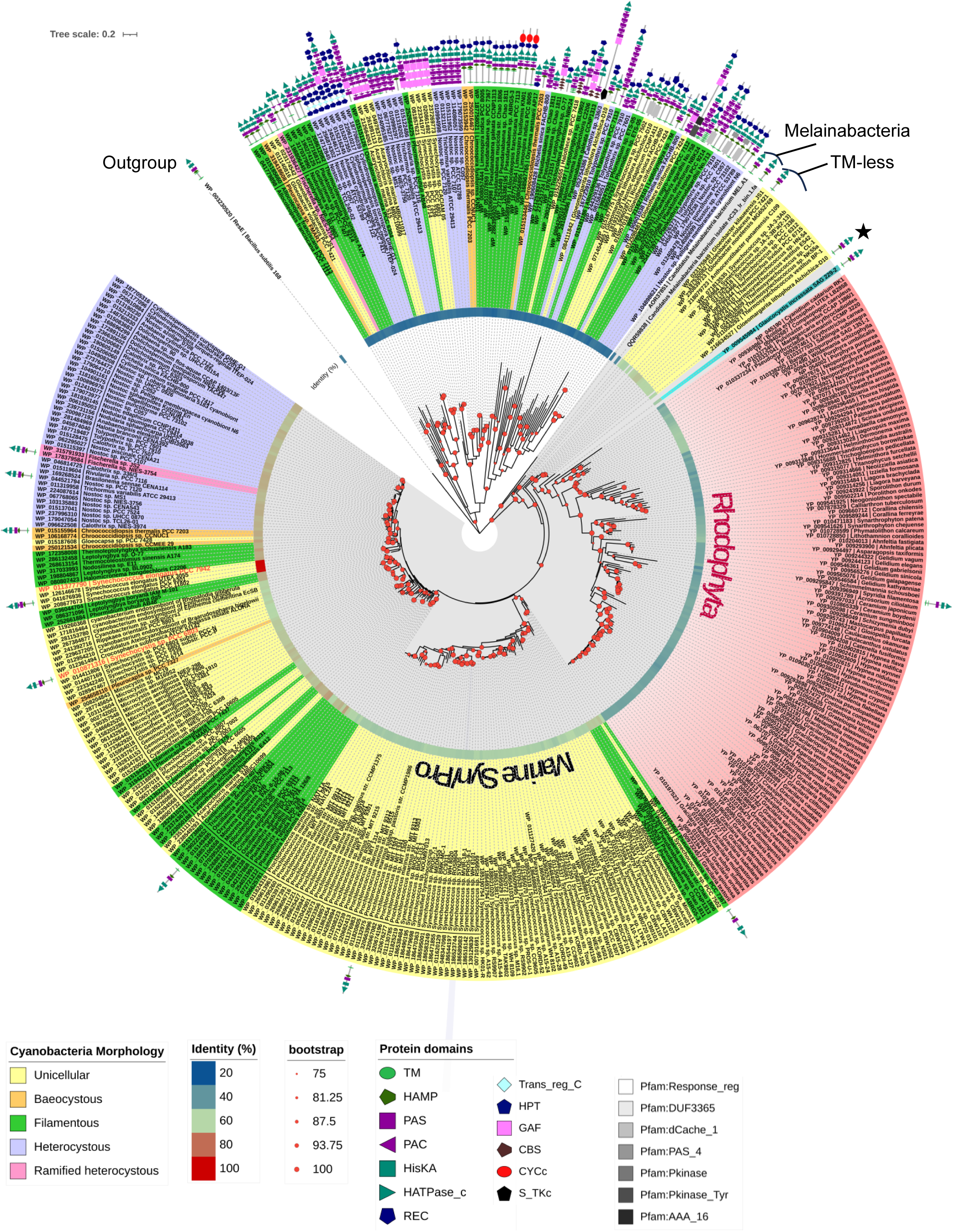
Phylogenetic tree of Hik33/NblS homologs and similar HKs. A phylogenetic tree was created by using maximum-likelihood method. Branches with white or grey background show other HKs clade or Hik33/NblS clade, respectively. Each node is labeled with refseq protein ID and organism name, colored according to the cell morphology of cyanobacteria. Eukaryotes, *Glaucocystis* and Rhodophyta are shown in blue and red respectively. TM-less indicates the specific clade of thylakoid membrane less cyanobacteria. ★ highlights the clade of *Gloeomargarita lithophora*. The entire clade of marine *Synechococcus* and *Prochlorococcus* is highlighted as Marine Syn/Pro. *Synechococcus* NblS and *Synechocystis* Hik33 are emphasized with red colored label. Bootstrap values more than 75% based on 1000 ultrafast replicates is shown with red filled circle. Amino acid sequence identity (%) against *Synechococcus* NblS is visualized with the heatmap. Protein domains annotated by SMART are illustrated for other HKs and representative Hik33/NblS orthologs.

## Discussion

The highly conserved histidine kinase in cyanobacteria, Hik33/NblS, is responsive to various unfavorable conditions, however, it has been elusive how this unique kinase is able to respond to such divergent stresses. In this study, we found that the histidine kinase NblS in *Synechococcus* responds to multiple abiotic stress conditions like Hik33 in *Synechocystis*, and these responses were alleviated in the presence of electron acceptor DCBQ which oxidize PSII (Fig 2). Cell fractionation analysis revealed the thylakoid membrane localization of NblS (Fig. 3B), and it was found that a dimeric NblS (hereafter NblS_2_) is associated with the PSII assembly intermediate RC47-like complex at least including D2 and CP47 proteins without CP43 (Fig. 4C). Together with other lines of evidence, we propose that NblS signaling is triggered by the changes in the redox state of Q_A_, the primary quinone acceptor of PSII. NblS/Hik33 proteins were found in a monophyletic clade in cyanobacteria apart from other HKs, and thus it was suggested that the underlying sensory mechanism is generally conserved.

It is of note that NblS response-inducing stresses result in the occurrence of the PSII photoinhibition. PSII photoinhibition is induced by excess light energy and is enhanced under various stress conditions (Nishiyama and Murata, 2014), where the PSII primary electron acceptor Q_A_ is over-reduced. Thus, we examined the NblS response in the presence of Q_B_ oxidizing reagent DCBQ and found that the response, as monitored by the *hliA* transcript accumulation, was alleviated under all conditions examined (Fig. 2). Especially, the Q_B_ site blocker DCMU induced the NblS response and DCBQ cancelled this activation. Therefore, these results strongly support the Q_A_ over-reduction model. Recently, involvement of the Q_A_ reduction in the Hik33 signaling in *Synechocystis* was also suggested based on the correlation between the response and the decrease of PSII quantum yield (Kato et al. 2022). However, the authors also suggested the involvement of PSI acceptor side for the signaling obscuring the sensory mechanism. Another study also proposed the photoreduction of the PQ pool and consequent Q_A_ reduction triggers the low temperature response in *Synechocystis* (Mironov et al. 2019). Our present study is consistent with those studies, and here we further postulate that the Q_A_ over-reduction but not membrane fluidity nor other abiotic stress itself is the sensory cue for the NblS/Hik33 kinases.

NblS was detected as two protein complexes by the CN-PAGE gel (Fig. 4A). These complexes lack CP43 and contain CP47 and D2 (Fig. 4C, S2C). This suggests that NblS binds to PSII-related subcomplex rather than oxygen-evolving PSII. Previous studies have shown that assembly of PSII involves a variety of subcomplexes with assembly factors (Komenda et al. 2024). So far, there are none of known complexes interacting with Hik33/NblS. Based on our results, we suggest that the complexes containing NblS_2_ are approximately 350 kDa and 700 kDa. This MW estimation is postulated because the position of the complexes in which NblS was detected coincides with the chlorophyll fluorescence emitted from the PSII monomer and PSII dimer (Fig. 4A). However, NblS_2_ is 150 kDa and the remaining MW of the lower band observed in the CN-PAGE gel is 200 kDa. Taking into account the known PSII-related subcomplexes, RC47 is the candidate of the partner of NblS. RC47 contains CP47 and D2 protein and does not contain CP43 and has MW 220 kDa (Boehm et al. 2012). If we consider the known MW of RC47, the MW of NblS_2_-RC47 would be 370 kDa, which is similar to the PSII monomer. Furthermore, RC47 has been reported to exist as a dimer. Therefore, if NblS_2_-RC47 becomes a dimer using RC47 as a scaffold (NblS_2_-RC47-RC47-NblS_2_), the MW is estimated to be 740 kDa, which is also similar to the MW of PSII dimers. However, this study has not identified all the proteins in the complex, which will be clarified in future studies. Thus currently, we propose that NblS interacts with RC47-like complex and then makes PSIIsos (Photosystem 2 stress <u>o</u>bserving scaffold) (Fig. 6).

**Figure 6.**
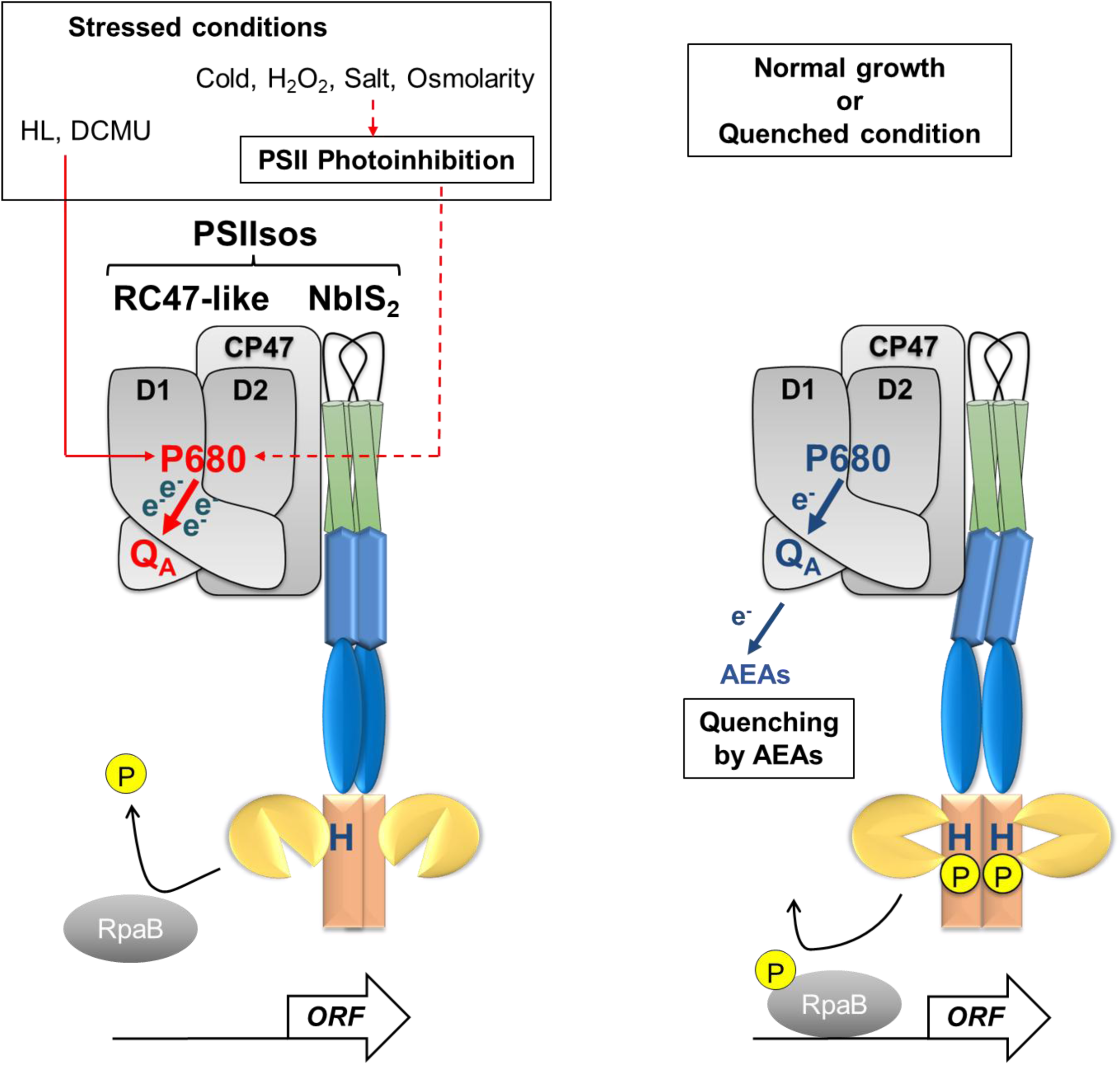
The schematic model of the NblS signaling under stressed conditions and quenching by PSII electron acceptors. NblS signaling is occurred in the protein complex PSIIsos (Photosystem 2 stress <u>o</u>bserving scaffold) composed by NblS_2_ and RC47-like. Under stressed conditions, the reaction center P680 and the primary quinone acceptor Q_A_ are reduced (colored with red) by the direct way (red line) or through the PSII photoinhibition (red broken line). The reduced Q_A_ causes NblS functional change from kinase to phosphatase, then RpaB is dephosphorylated and released from the promoter region of regulon genes shown as *ORF*. By addition of alternative electron acceptors (AEAs) as well as under normal condition, P680 and Q_A_ are oxidized (colored with blue). It leads the maintaining of NblS kinase activity and RpaB is then phosphorylated and bound to DNA subsequently. NblS is described with the secondary structure as in Figure 3A. e^-^ indicates electrons in PET. Phosphates involved in the phospho-transfer events are shown as P in yellow.

We demonstrated that NblS signaling is triggered by sensing the reduction of Q_A_ within RC47-like complex via direct interaction. How is the redox status of RC47-like complex sensed by NblS in PSIIsos? It’s possible that NblS binds a quinone molecule to receive electrons from RC47-like complex, however NblS has neither quinone binding motifs nor redox active cysteine residues in its amino acid sequence as reported in other HKs (Franza and Gaudu, 2022; Ibrahim et al. 2020). Current reasonable hypothesis is that the signal transduction from RC47-like complex to NblS might be mediated via protein conformational rearrangements. Our prediction of the NblS_2_-CP47-D2 structure showed that NblS_2_ interacts with CP47 via a transmembrane region (Fig. S3B). It is noteworthy that these amino acids of CP47 in the interaction site are necessary for photoautotrophy (Clarke et al. 2002). In the agreement with both *in vivo* and *in vitro* previous research, transmembrane region and HAMP domain are important for switching its function as phosphatase, while the lumenal loop in between transmembrane regions is dispensable (Bartsevich and Shestakov, 1995; Leusenko et al. 2024; López-Redondo et al. 2010; Shimura et al. 2012). Thus, the conformational change of CP47 upon the reduction of Q_A_ might lead to the rotation of HAMP domain. The rotation of the HAMP domain is known to switch the kinase activity or phosphatase activity of HKs (Bhate et al. 2015). Additionally, we aligned the modeled NblS_2_-CP47-D2 structure with the recently revealed structure of Psb28-RC47 (PDB: 7DXA) (Xiao et al. 2021). The position of NblS_2_ overlapped with Psb34 and UTP (Fig. S4). It has been suggested that the Psb34 binding site is the same as the Hlips binding site of RC47 in *Synechocystis* (Rahimzadeh-Karvansara et al. 2022). Since *hli* genes encoding Hlips are regulated by Hik33/NblS in both *Synechocystis* and *Synechococcus*, it is rational that Hk33/NblS senses stress at the same site as Hlips and makes the feedback loop for acclimatation via the expression of *hli* genes. Further study will unveil the whole mechanism from the sensing to acclimation against stressed state of photosynthesis by Hik33/NblS.

Together with our data of the NblS protein complex, our phylogenetic analysis highlights the evolutionarily vital role of Hik33/NblS/Ycf26 since the emergence of oxygenic phototrophs (Fig. 5). However, the conservation of Ycf26 in eukaryotes is not simple, even though PSII is fully conserved. For instance, *ycf26* is encoded in the chloroplast genome of *Glaucocystis incrassata* but not in the other Glaucophyta and lost completely in the green lineage. In Rhodophyta, the conservation is mosaic as that many species have intact Ycf26 but some species do not have and some species have Ycf26 truncated from its N-terminus region (Supplementary Dataset S3). Furthermore, not only primary symbiotic algae but also some of secondary symbiotic algae keep *ycf26* in their chloroplast genome (Duplessis et al. 2007), however the function of Ycf26 in chloroplast is still elusive. In green lineage, sensory kinases working in chloroplast are encoded in nuclear genome while there are no known kinases with PSII associated sensing mechanisms (Bellaflore et al. 2005; Bonardi et al. 2005; Puthiyaveetil et al. 2008). Thus, Hik33/NblS/Ycf26 can be a good marker for the evolution of cyanobacteria and chloroplast, on the other hand, it’s also target of rewiring the regulatory mechanism of chloroplast gene expression. Future research of Ycf26 function in eukaryotic chloroplast and algal nuclear genomes will unveil the evolutionary change of plastid transcriptional regulation.

## Materials and Methods

### Growth condition of cyanobacteria

*Synechococcus elongatus* PCC 7942 was grown by BG-11 media buffered with 20 mM HEPES-KOH pH 8.0 as 1.5% agar containing plates or liquid under continuous white fluorescent lamp 20 μmol m^-2^ s^-1^ at 30°C. Liquid culture was supplemented with 2% CO_2_ bubbling. For selection of mutants, kanamycin and spectinomycin were used as 10 µg/ml and 40 µg/ml, respectively.

### Strain construction

For NF strain, the ORF of *nblS* was amprified from *Synechococcus* WT genome by PCR with the primer sets (Supplementary Table S1) and cloned into SacI/BamHI site of pKS-hik2-FLAG to obtain c-terminal FLAG-tagged *nblS* (Hasegawa et al. 2020). The *nblS*-FLAG fragment was excised by SacI/SmaI treatment and inserted to SacI/SalI site of pSB (Hasegawa et al. 2020) with SmaI/SalI treated 928 bp of *nblS* downstream fragment. First selection by kanamycin and secondary counter selection by sucrose were done as previously (Hasegawa et al. 2020).

For NF43H strain, the 736 bp fragment of *psbC* from its 3’-end without the stop codon and the 729 bp fragment of *psbC* downstream region were amplified and cloned to SacI/SmaI and HincII site of pBΩH, respectively (Hasegawa et al. 2020). The plasmid was used for transformation of NF strain and selected by spectinomycin.

RBF strain was constructed as previously reported (Hanaoka and Tanaka, 2008).

### Protein analysis by SDS-PAGE, CN-PAGE and immunoblot

Cells from 10 mL liquid culture of NF strain at OD_750_ = 0.4-0.6 were collected by centrifugation (7,000 × *g*, 2 min, 4°C). Cells were resuspended in 200 µL of Buffer 1 (50 mM HEPES-NaOH pH 7.0, 5 mM CaCl_2_, 10 mM MgCl_2_, 25 % w/v glycerol) and broken with equal volume of grass beads (G4649, Sigma-Aldrich) by vortexing at maximum speed (1 min, 1 min interval on ice, 10 cycles, 4°C). After the cell disruption, unbroken cells and cell debris were removed by centrifugation (3,000 × *g*, 2 min, 4° supernatant was sedimented by centrifugation (18,800 × *g*, 20 min, 4°C). The pellet was resuspended by Buffer 1 and the aliquot was used for chlorophyll extraction. Chlorophyll concentration was calculated after extraction by 100% Methanol, as previously (Porra et al. 1989). Samples were denatured with LDS sample buffer and conducted to Tris/Mes SDS-PAGE with 6 M urea containing 10% polyacrylamide gel (Kashino et al. 2001).

For CN-PAGE, cells from 200 ml culture were collected as mentioned above and suspended in 20 mL of Buffer 1. One half volume of zirconia/silica beads (11079101z, Biospec) were added and cells were broken by Multibeads Shocker (YASUI KIKAI) at 1,700 rpm (30 sec, 2 min interval on ice, 10 cycles). After cell disruption, thylakoid membrane was separated as mentioned above and resuspended at 1 mg chlorophyll/ml in Buffer 1. The suspension was supplemented with equal volume of 2% *n*-Dodecyl-α-D-Maltoside (hereafter α-DDM, AG-CC1-0006-G001, KOM) in H_2_O and incubated (5 min, on ice, dark). After removing insoluble parts by centrifugation (18,800 × *g*, 5 min, 4°C), the sample corresponding to 5 µg chlorophyll were separated by CN-PAGE with 4-13% gradient polyacrylamide gel at (4 mA, 4°C, 180 min) as described previously (Koskela et al. 2020). After electrophoresis, chlorophyll fluorescence from the gel was observed by Cyanoview and Luminograph (ATTO) with a band pass filter SC-68 (Fujifilm). For immunoblot, the gel was incubated by Buffer 2 (25 mM Tris, 192 mM Glycine, 0.1% SDS, 20% Methanol) for 15 min before blotting. NblS-FLAG, CP43-His and D2 were detected by anti-FLAG antibody (1E6, Fujifilm Wako), anti-His antibody (D291-3, MBL) and anti-D2 antibody (AS06-146, Agrisera) respectively. CP47 and PsaAB were detected by each specific antibody.

### Cell fractionation by Sucrose density centrifugation

Cell fractionation was done base on the previous report with modifications as followed (Schottkowski et al. 2009). 100 mL culture of NF strain at OD_750_ = 0.4-0.6 was harvested by centrifugation (7,000 × *g*, 5 min, 4°C). The cell pellet was once washed by 25 mM HEPES-NaOH pH 7.0 and resuspended in 1 mL of Buffer 3 (25mM HEPES-NaOH pH 7.0, 5 mM EDTA, 10 mM NaCl, 0.6 M sucrose and 1 mM phenylmethylsulfonyl fluoride). One half volume of zirconia/silica beads were added and cells were broken by vortexing at the highest speed (30 sec, 2 min interval on ice, 10 cycles, 4° centrifugation (3,000 × *g*, 2 min, 4°C), the supernatant was collected and incubated with 1 µL of Recombinant DNase I (RNase-free) (TaKaRa) under dark (15 min, on ice). 395 µL of the sample was mixed with 292 µL of 90% sucrose in a 2.2 mL Open-Top Thinwall Ultra-Clear Tube (347356, Beckman Coulter). Then, the mixture was overlaid with 412 µL of 39% sucrose, 550 µL of 30% sucrose and 687 µL of 10% sucrose prepared in Buffer 3. After centrifugation (77,000 × *g*, 15 hours, 4° bucket rotor (TLS-55, Beckman Coulter), the gradient was fractionated into Fraction I-V as mentioned in the previous report (Schottkowski et al. 2009). In case of further analysis by SDS-PAGE and immunoblot, proteins from each fraction were precipitated and concentrated by trichloroacetic acid as previously reported (Nimura et al. 2001).

For further separation, 176 µL of Fraction V was diluted with 264 µL of 5 mM HEPES-NaOH pH 7.0 to 20% sucrose and overlaid to 1.76 mL of 30-60% linear sucrose gradient. After centrifugation (77,000 × *g*, 15 hours, 4 °C), the gradient was fractionated into Fraction 1-14 by taking 140 µL from top to bottom of the tube. All fractions were used for SDS-PAGE in equal amounts.

### Protein crosslink, immunoprecipitation and His_6_-tag purification of PSII

4 L culture of NF43H strain at OD_750_ = 0.4-0.6 were harvested by centrifugation (5000 × *g,* 5min, room temperature) and once washed by 50 mM HEPES-NaOH pH 7.0. Cells were suspended in 40 mL of Buffer 1 and supplemented with 200 mM DSP stock solution at final concentration 5 mM. As the protein cross-link reaction, cell suspension was incubated in dark (30 min, room temperature, with gentle rotation) and quenched by adding 1 M Tris-HCl pH 7.5 at final concentration 60 mM under dark (15 min, room temperature, with gentle rotation). Cells were collected by centrifugation (7,000 × *g*, 2 min, room temperature), once washed by Buffer 1 and kept in −80° disruption and preparation of solubilized thylakoid membrane were done by same procedure in CN-PAGE analysis as mentioned above. Solubilized thylakoid membrane corresponding 1 mg chlorophyll were used for FLAG immunoprecipitation and His-tag purification as followed.

For FLAG immunoprecipitation, 3.0 mg of Dynabeads Protein G (10004D, Invitrogen) equilibrated by Buffer 1 with 0.03% α-DDM was added to the solubilized thylakoid membrane solution. In addition, 10 µg of anti-FLAG antibody (1E6, Fujifilm Wako) was supplemented and incubated under dark (120°C, with gentle rotation). As the negative control, 10 µg of mouse IgG (SC-2025, Santa Cruz Biotechnology) was used. After incubation, the Dynabeads was collected by centrifugation (500 × *g*, 2 C min, 4°C) and washed three times by 5 volume of Buffer 1 with 0.03% α-DDM. Finally, the Dyanabeads was denatured by equal volume of 2 x LDS sample buffer, to cleave protein cross-link, incubated with 20 mM DTT (30 min, 37°C). For protein analysis, SDS-PAGE and immunoblot were done as mentioned above.

For His-tag purification, 1/10 bed volume of cOmplete His-tag Purification Resin (Roche) equilibrated by Buffer 1 with 0.03 % α-DDM was added to the solubilized thylakoid membrane solution and incubated under dark (120 min, 4°C, with gentle rotation). After incubation, the resin was washed 10 times with 1 mL of Buffer 1 containing 0.03 % α-DDM and used for protein analysis as mentioned in FLAG immunoprecipitation.

### Stress treatments and RNA analysis

Liquid culture of *Synechococcus* WT at OD_750_ = 0.4-0.6 was used for stress treatments as followed. HL treatment was done by shedding 120 μmol m^-2^ s^-1^ white fluorescent lamp. DCMU (D2425, Sigma-Aldrich), NaCl and Sorbitol were added at 10 µM, 0.5 M and 0.5 M respectively. For Cold treatment, the culture was transferred to a 14°C -water bath. For Mock treatment, ethanol was added at final 0.1%. BQ (171-00242, Fujifilm Wako), DCBQ (D0344, Tokyo Chemical Industry), DMBQ (428566, Sigma-Aldrich) and DQ (T0672, Tokyo Chemical Industry) were prepared at 1 mM in ethanol and used at final concentration 100 µM. Silicomorybdate (708348, Sigma-Aldrich) were used at final concentration 0.2 mg/ml or 2 mg/ml. For sampling, these alternative electron acceptors were added and incubated for 5 min prior to stress treatment. All stress treatment were done for 30 min and cells were transferred on ice. Collection of cells was done by centrifugation (7,000 × *g*, 2 min, 4°C) and stored at - 80°C until RNA extraction

Extraction of total RNA from *Synechococcus* and Northern blot analysis were done as previously reported (Kobayashi et al. 2017). The DNA probes of *hliA*, *rnpB* and *psaB* were prepared by the primers shown in Supplementary Table S1. 3 µg of total RNA was applied to Northern blot analysis.

### Chromatin immunoprecipitation-qPCR

ChIP-qPCR of RBF strain was done as the previous report (Hanaoka and Tanaka, 2008). Primers used for qPCR are shown in Supplementary Table S1.

### Phylogenetic analysis

For Cyanobacteria/Melainabacteria group, genome and protein sequence data from the assembly level as chromosome and complete was downloaded from NCBI assembly in 20231226. For eukaryotic plastids, RefSeq sequence data from 15,000 species was obtained from NCBI datasets Organelle in 20231226. Thus, we constructed our database containing 308 cyanobacteria, 3 Melainabacteria and 15,000 eukaryotic plastids. ResE of Bacillus subtilis 168 (RefSeqID: WP_003230520.1) was used as the out group as previously mentioned (Los et al. 2010). Homologous proteins to NblS of *S. elongatus* PCC 7942 (RefSeqID: WP_011377790.1) was searched by BLASTP with the query sequence (NblS 213-638). NblS orthologs were defined according to the E-value (up to 1.00E-65 in Cyanobacteria/Melainabacteria and 1.00E-10 in plastids, respectively). In case of other HKs, the result of BLASTP was restricted with query coverage more than 90%. After identical sequences were removed manually, non-redundant protein sequences were aligned by MAFFT version 7.490 with default parameters and trimmed by trimal with “auto” (Capella-Gutiérrez et al. 2009; Katoh and Standley, 2013). After trimming of the alignment, redundant sequences were manually removed again. The alignment was used to construct a maximum likelihood phylogenetic tree by IQ-Tree version 2.1.2 in CIPRES Science Gateway with LG+I+G model and the Gamma shape parameter 4 (Miller et al. 2010; Minh et al. 2020). The phylogenetic tree was annotated and visualized by iTOL v7 web service (Letunic and Bork, 2024). Same procedure was applied to create a tree for only NblS orthologs with full length of their amino acid sequences.

### Structure modelling

Amino acid sequences of NblS, CP47 and D2 of *S. elongatus* PCC 7942 were obtained from Uniprot (NblS: Q8RQ68, CP47: P31094 and D2: P11005). Structure modelling was done by Alphafold3 in AlphaFold Server. Visualization of the modelled structure and the distance analysis of lysine residues were done by UCSF ChimeraX version 1.7.1 (Meng et al. 2023). Then, the modelled structure was aligned with Psb28-RC47 (PDB: 7DXA) as the reference structure by Matchmaker function of UCSF ChimeraX with “Best-aligning pair of chains between reference and match structure”.

## Acknowledgments

We thank Dr. Masahiko Ikeuchi and Dr. Yuichiro Takahashi for gifting antibodies against D1 and CP1 respectively, Dr. Ryouichi Tanaka and Dr. Atsushi Takabayashi for teaching the techniques of CN-PAGE, Mr. Yuta Kobari for the preliminary test of cell fractionation method. We also thank Dr. Josef Komenda, Dr. Jürgen Tomasch and Dr. Ondřej Prášil for suggestions and critical reading of the paper.

## Author contributions

T.T. and K.T. designed the study. T.T. performed the experiments, analyzed the data and wrote the manuscript. K.T. supervised the project and wrote the manuscript.

