## Supplementary figure 1-5 for "An Evolutionary Conserved Multi-Stress Sensory Histidine Kinase NblS Associates With Photosystem II Proteins And Responds To Its Redox Status In The Cyanobacterium *Synechococcus elongatus* PCC 7942"

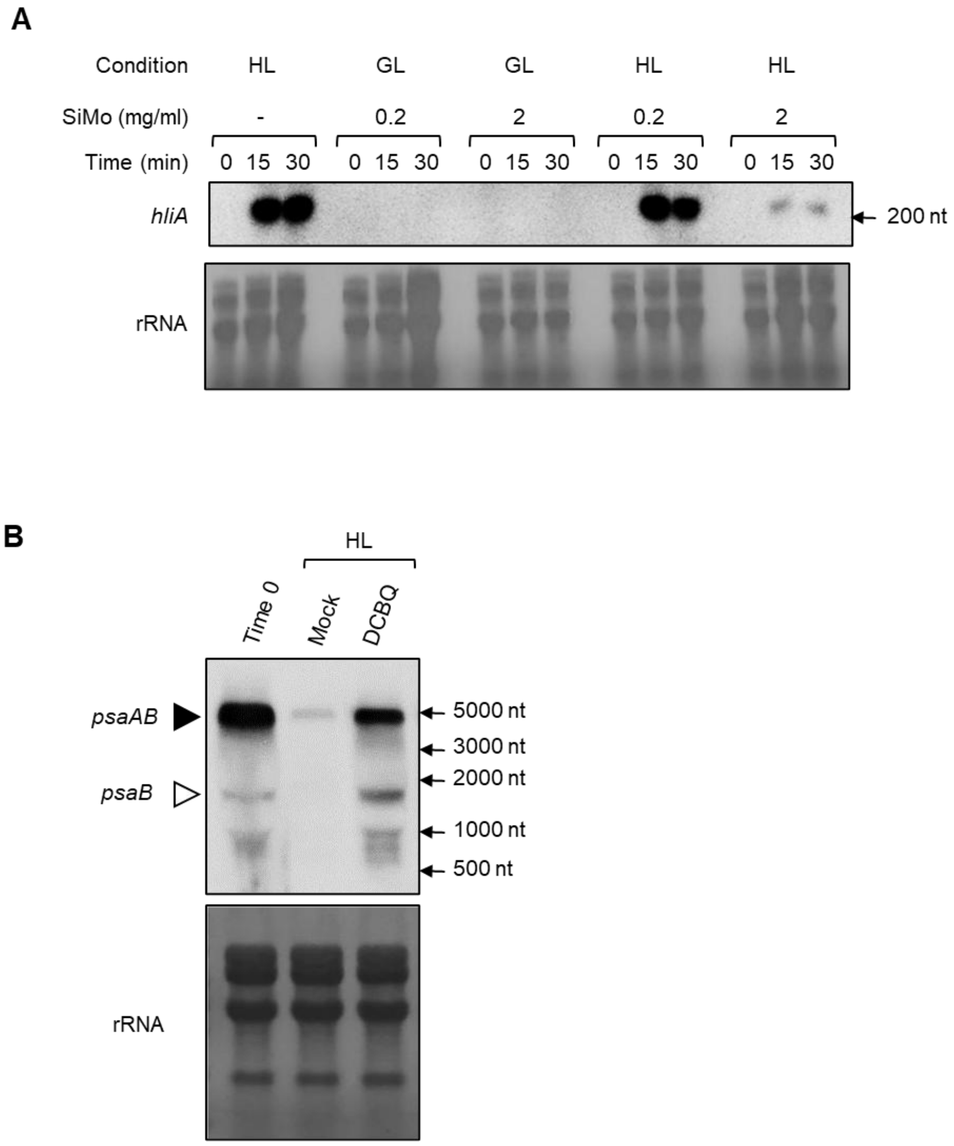


**Figure S1.** Quenching analysis of HL stress response. (**A**) The result of Northern blot against *hliA* transcript with non-quinone derivative PSII alternative electron acceptor silicomolybdate (SiMo). GL and HL indicate growth light and high light shed on *Synechococcus* WT culture respectively. SiMo with indicated concentration was added to the culture and incubated for 5 min before each light treatment. (**B**) The result of Northern blot against *psaAB* transcript with DCBQ under HL. Mock and DCBQ indicate the samples treated with 0.1% ethanol or 100 µM DCBQ for 5 min before transferring to HL condition respectively. Time 0 indicates the sample before HL treatments. Specific probe to *psaB* was used for the detection. Black arrowhead and white arrowhead correspond to *psaAB* and *psaB* according to the size marker respectively. rRNA is the image of Methylene Blue stained membrane as the loading control. 3 µg of total RNA were loaded in each lane.

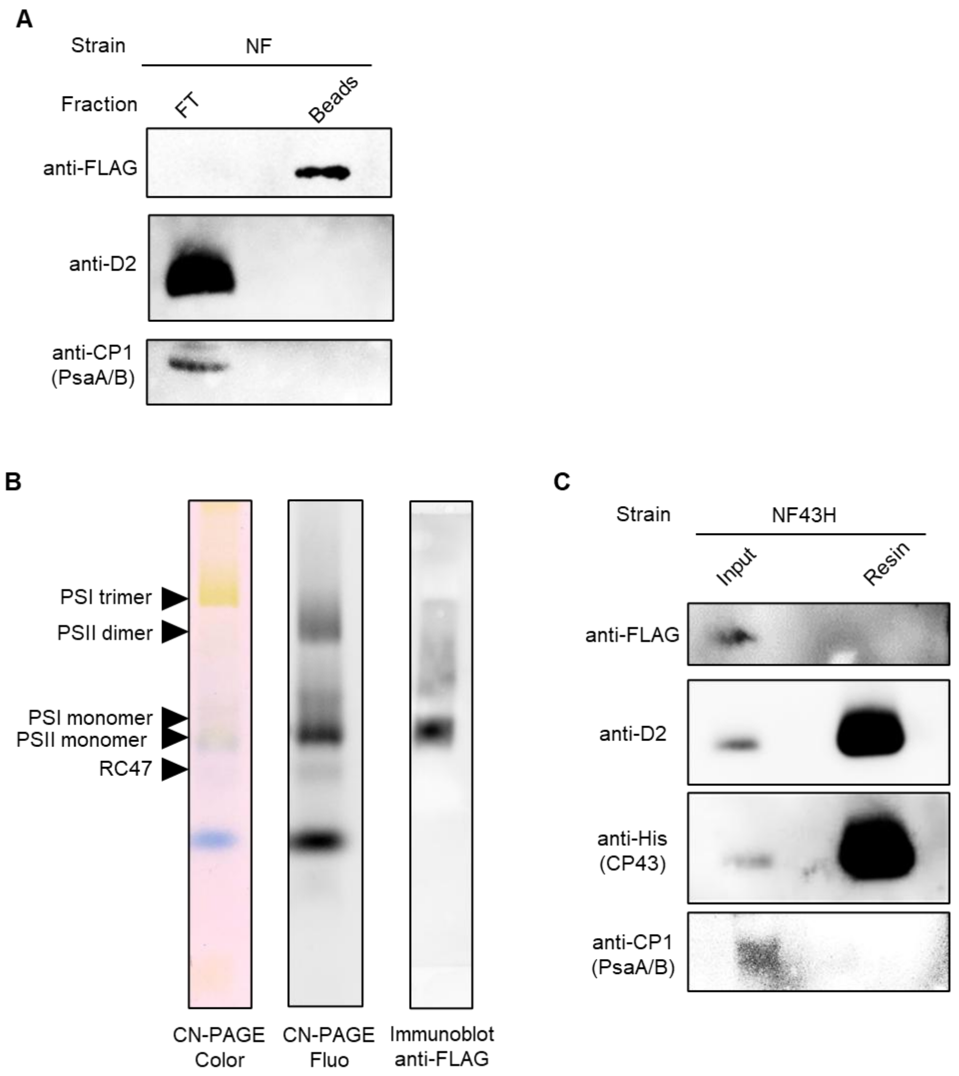


**Figure S2.** Analysis of the NblS protein complex with and without cross-link. (**A**) The result of immunoblot from the FLAG pull-downed thylakoid membrane of NF strain without protein cross-link. Flow through (FT) and Dynabeads (Beads) after immunoprecipitation were used for SDS-PAGE and immunoblot. (**B**) The result of CN-PAGE from DSP cross-linked thylakoid membrane of NF43H strain. CN-PAGE Color is the RGB gel picture, CN-PAGE Fluo is the chlorophyll fluorescence gel image excited at 505±25 nm and detected with a red filter. The positions corresponding to each photosystem complexes were estimated from the previous work (Koskela et al. 2020). The position of NblS-FLAG detected by immunoblot is shown in Immunoblot anti-FLAG. (**C**) The result of immunoblot after the His-tag purification of PSII. DSP cross-linked thylakoid membrane of NF43H strain was used. SDS-PAGE was done after cleavage of DSP cross-link by DTT. Input and Resin indicate the thylakoid membrane corresponding to 0.5 µg chlorophyll before purification and $Ni^{2+}$ resin after purification.

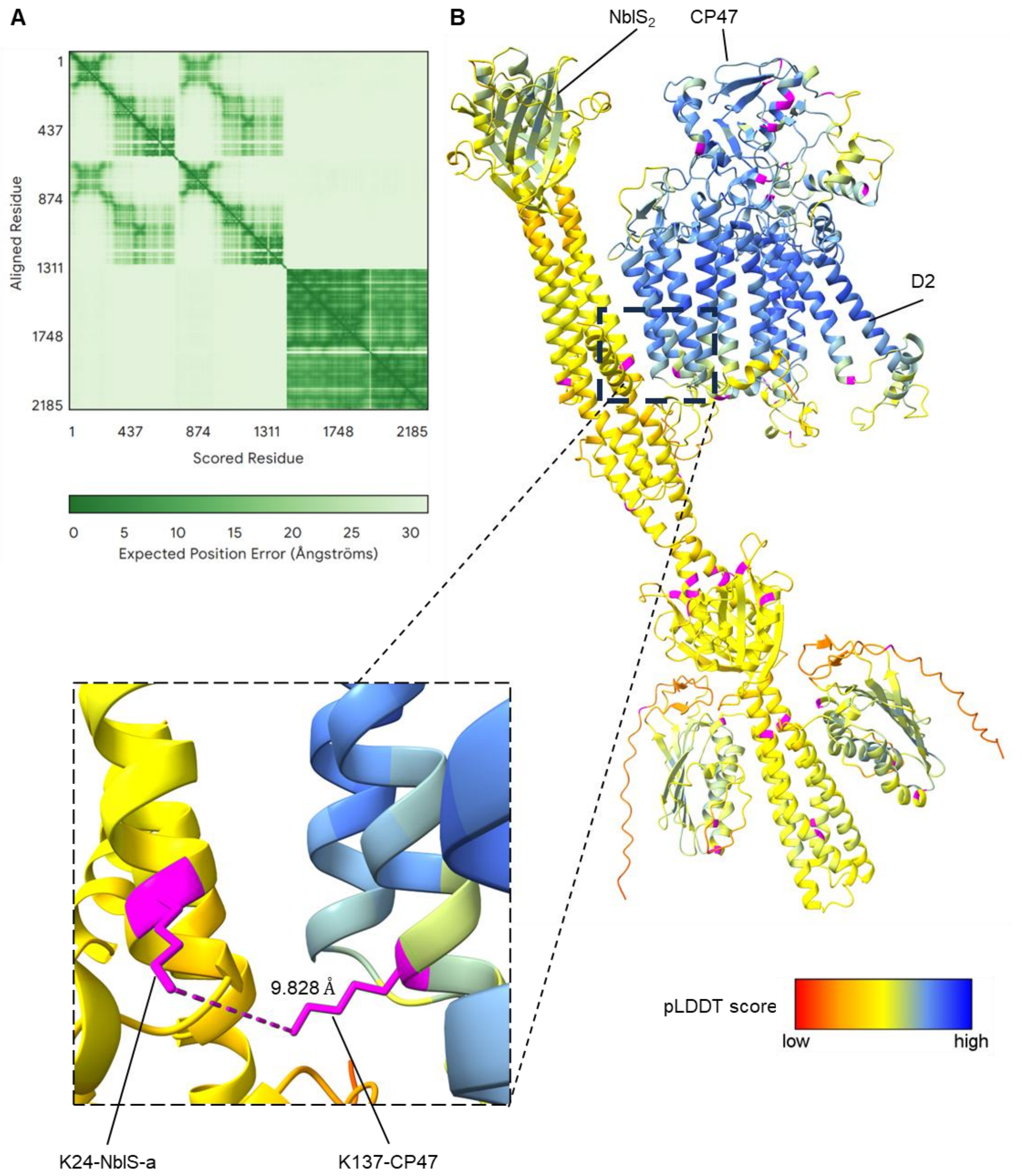


**Figure S3.** Modelling of the NblS$_2$-CP47-D2 complex by Alphafold3. (**A**) The image of expected position error plot is shown (residue positions as followed, NblS-a: 1-664, NblS-b: 665-1328, CP47: 1329-1836 and D2: 1837-2188). (**B**) The structure of modelled NblS$_2$-CP47-D2. Except for lysine residues, amino acids in the structure are colored by pLDDT score as represented in the heatmap. To search the position of DSP cross-link, lysine residues (K) are highlighted with magenta. The region of K24 from NblS-a and K137 from CP47, which are the candidates of DSP cross-link, is enlarged in the panel with broken lines. The distance between primary amine side chains of these lysine residues is also shown (9.828 Å).

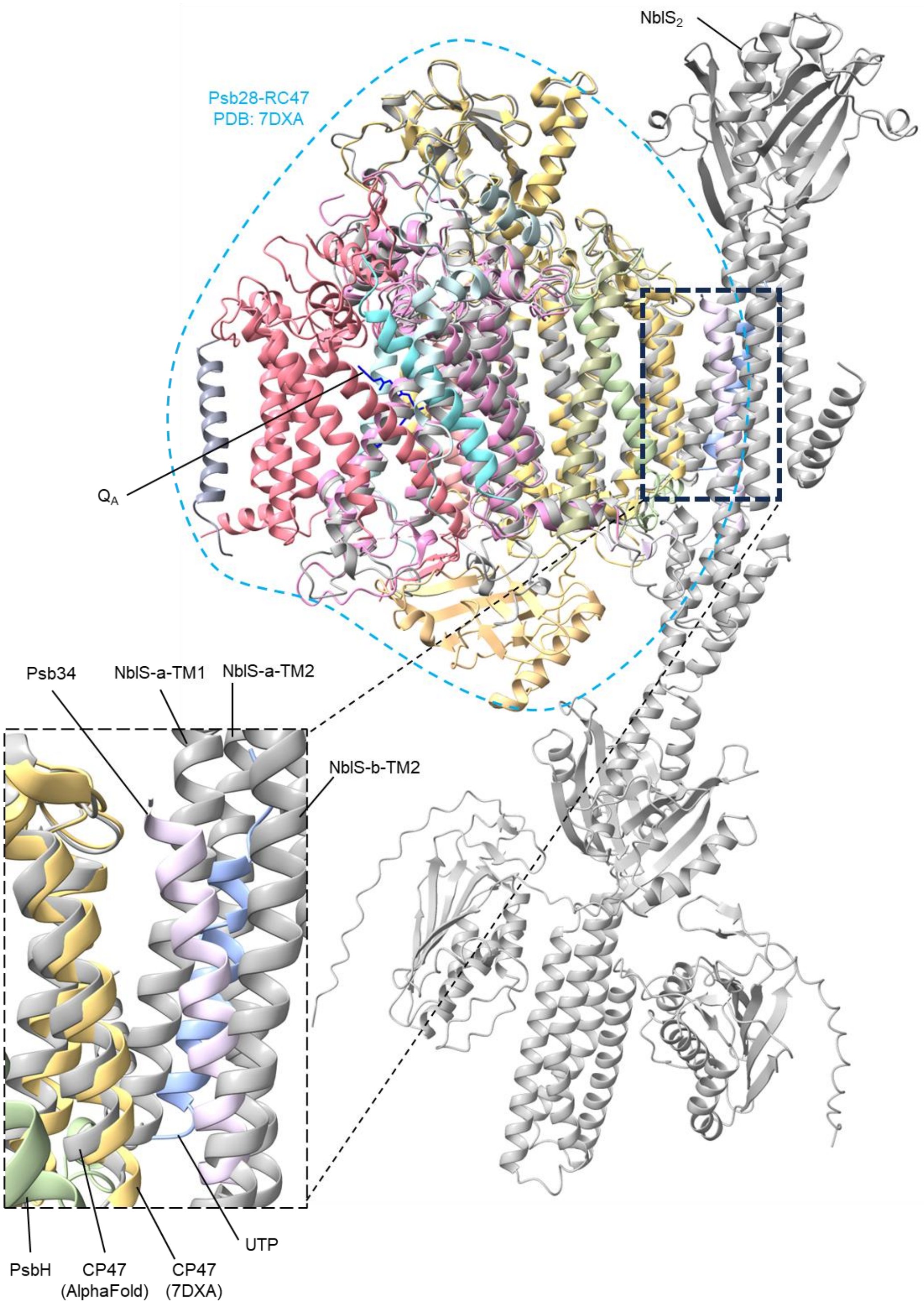


**Figure S4.** Alignment of the NblS$_2$-CP47-D2 modelled by Alphafold3 and published Psb28-RC47 structure. The modelled NblS$_2$-CP47-D2 structure (gray) was aligned with Psb28-RC47 (PDB: 7DXA). Psb28-RC47 is highlighted with light bule broken line. The $Q_A$ molecule is colored with dark blue. The putative interaction site of NblS$_2$ is enlarged in the panel with broken lines. In the panel, NblS-a-TM1 and NblS-b-TM2 correspond to the transmembrane regions from each chain as shown in Fig. 3A.

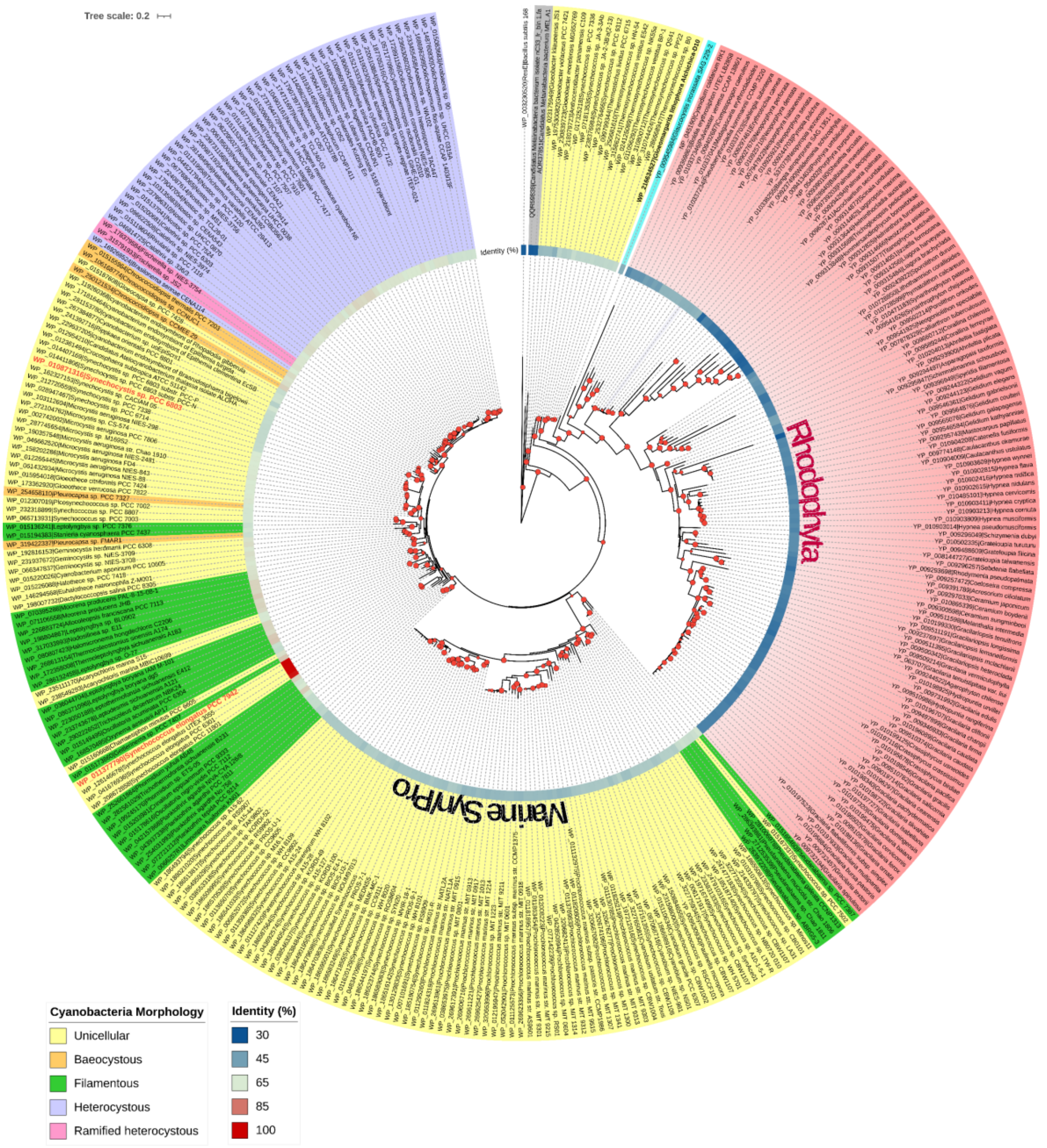


**Figure S5.** Maximum-likelihood tree created with full length of Hik33/NblS orthologs. Each node is labeled with refseq protein ID and organism name, colored according to the cell morphology of cyanobacteria. Eukaryotes, *Glaucocystis* and Rhodophyta are shown in blue and red respectively. The entire clade of marine *Synechococcus* and *Prochlorococcus* is highlighted as Marine Syn/Pro. *Synechococcus* NblS and *Synechocystis* Hik33 are emphasized with red colored label. Bootstrap values more than 75 % based on 1000 ultrafast replicates is shown with red filled circle. Amino acid sequence identity (%) against *Synechococcus* NblS is visualized with the heatmap.
